# A closer look at the timecourse of mind wandering: pupillary responses and behaviour

**DOI:** 10.1101/869768

**Authors:** Claudia Pelagatti, Paola Binda, Manila Vannucci

## Abstract

Mind wandering (MW) refers to the shift of attention away from a primary task and/or external environment towards thoughts unrelated to the task. Recent evidence has shown that pupillometry can be used as an objective marker of the onset and maintenance of externally-driven MW episodes. In the present study we aimed to further investigate pupillary changes associated with the onset and duration of self-reported MW episodes.

We used a modified version of the joint behavioural-pupillometry paradigm we recently introduced. Participants were asked to perform a monotonous vigilance task which was intermixed with task-irrelevant cue-phrases (visually presented verbal cues); they were instructed to interrupt the task whenever a thought came to mind (self-caught method) and to indicate the trigger of their thought, if any. We found systematic pupil dilation after the presentation of verbal cues reported to have triggered MW, compared with other verbal cues presented during a supposedly on-task period (i.e., the period immediately following the resuming of the task after a self-caught interruption and MW report). These results <u>confirm that pupil diameter is sensitive to the changes associated with the onset of MW and its unfolding over time</u>.

Moreover, by computing the latency between the trigger presentation and the task interruption (self-catch), we could also estimate the duration of MW episodes triggered by verbal cues. However, a high variability was found, implying very large inter-event variability, which could not be explained by any of the MW properties we acquired (including: temporal focus, specificity, emotional valence). Our behavioural and pupillometry findings stress the need for objective measures about the temporal unfolding of MW (while most studies focus on arbitrary time-window preceding self-reports of MW).

## Introduction

In our daily lives, we may notice that our attention drifts away from the ongoing task and external environment, and our mind starts wandering elsewhere, towards task-unrelated private thoughts and feelings such as memories, future plans, current concerns. This *“shift in the focus of attention away from the here and now towards one’s private thoughts and feelings”* [1, p. 818], which mainly occurs spontaneously, is referred to as mind wandering (hereafter MW; for review, 2).

The phenomenon of MW is pervasive, being more likely and frequent during automatized and well-practiced tasks (e.g., driving, reading). Our understanding of the neurocognitive process of MW has dramatically increased over the past two decades, and great benefits arise from the use of the “strategy of triangulation” [2] whereby self-reports, behavioral measures and physiological measures are combined in the same study to make inferences about covert mental experiences.

During the last years, increasing attention has been paid to the investigation of the processes and events that prompt the initial occurrence of MW (its onset) and support its unfolding over time (process-occurrence framework, 3). Indeed, the possibility of empirically addressing this issue is strongly related to the identification of the triggers of MW: MW states need to be causally linked to preceding events in order to study the onset of these experiences and therefore to track their timecourse.

According to Klinger’s *current concerns* hypothesis [4,5,6] “spontaneous thoughts are probably *triggered* by cues (meaningful stimuli) that may be external in the environment or internal in the person’s own mental activity and that are associated with one or another of the individual’s goals” [6, p. 216]. Over the last years, an increasing number of studies [7–13] have investigated the potential contribution of environmental stimuli (not only goal-related stimuli) to MW. The results converge in showing that both task-relevant [7–9] and task-irrelevant external stimuli [10,11,13] might indeed act as triggers for MW, challenging the traditional view of MW as completely stimulus-independent [14] and self-generated mental activity [3].

At the methodological level, an important contribution has been provided by a new experimental paradigm, developed by Schlagman and Kvavilashvili (2008) and extensively used to investigate involuntary autobiographical memories [e.g., 16–20], and recently adapted to study MW [11,13].

In this paradigm, participants are presented with a long sequence of trials of mostly horizontal lines and have to detect an infrequent target (i.e., vertical lines). Participants are also exposed to cue-phrases presented in the centre of the screen (e.g., ‘missed opportunity’ or ‘wall mirror’), which they are told are irrelevant to the task. The experience of MW can be assessed during the task by using the self-caught method (i.e., instructing participants to stop the task every time they catch their mind not on-task/mind wandering) or the probe-caught method (i.e., interrupting participants during the task and asking them about their experience just immediately prior to the probe). With both methods, participants are asked to briefly describe their thoughts and to specify the trigger, if any.

The results obtained with this vigilance task revealed that (i) the majority of recorded MW episodes are reported as being triggered by the task-irrelevant verbal cues, (ii) the exposure to verbal cues may affect the temporal orientation of MW, increasing the proportion of past-oriented MW [11,13]. However, one of the limitations of these studies is that MW triggers are identified by introspective measures, i.e. asking participants to report what had triggered each MW event. In addition, the use of self-report measures does not allow the monitoring and tracking of the dynamics of MW, such as the changes in the attentional state associated with the onset and maintenance of MW.

In a first attempt to overcome these limitations, we combined self-report measures of MW and pupillometry [21]. Participants performed the vigilance task with task-irrelevant cue-words and MW was assessed by using the probe-caught method – interrupting the task and asking participants to report their thoughts and what had triggered them.

The results showed significantly larger pupil dilation following verbal cues reported as triggers of a MW episode compared to the control conditions. The pupil dilation increased over time, suggesting that a change in pupil diameter follows the onset of MW and accompanies its unfolding and maintenance over time. Given the well-known association between pupil dilation and cognitive or emotional load [22,23], we explained the pupil dilation observed after the onset of a MW episode in terms of the increased cognitive and emotional processing associated with the onset of MW and its unfolding, compared to the monotonous vigilance task.

In the present study, we aimed to replicate these findings within a similar paradigm (vigilance task with verbal cues) but using the self-caught rather than probe-caught procedure to assess the occurrence of MW episodes. In the self-caught method, participants were instructed to interrupt the task whenever they became aware of any task-unrelated mental contents, to report them and their triggers, if any. With this procedure, participants necessarily reported only thoughts that they were aware of, while they failed to report MW episodes that remained below the awareness threshold (zoning out, see 24). Previous studies have focused on unaware MW episodes, which are associated with poorer performance [e.g., 25] and with a more pronounced activation of the default mode and executive networks [26] compared to aware MW episodes. Thus, using the self-caught rather than the probe-caught method used in [21], here we focus on a more homogeneous subset of MW episodes (aware MW), to replicate and extend previous findings on the pupillary correlates of the dynamics of MW.

The combination of the behavioral paradigm with cue-words and self-caught method also allowed for addressing an additional question, about the duration of MW. Specifically, for those MW episodes that participants reported as being triggered by verbal cues shown on the screen, we could calculate the latency data or retrieval times, that is the time-interval between the presentation of the trigger and the MW report. This interval reflects the time needed for the arising of thoughts and the awareness of them. In the research field of MW, with a few exceptions [5,27,28] the question of the potential duration of MW has not been empirically addressed. Indeed, several studies have examined measures associated with MW states (i.e., reaction time variability, eye movements, fMRI BOLD signal) by using arbitrary time-windows length before self-reports of MW states, assuming that MW episodes occurred precisely in those windows and lasted for that period of time. Here we used participants’ self-reports to directly measure the latency of MW episodes triggered by cue-words; we also analyzed the association between latency and some of the main phenomenological properties of MW episodes (i.e., temporal focus, specificity, emotional valence), to explore the possibility that these variables regulate the temporal unfolding of MW.

## Materials and Methods

### Participants

Twenty-eight undergraduate students from the University of Florence (age range 19-32 years, M = 21.61 years, SD = 3.06 years, 16 females) volunteered to participate in the study. Four were excluded due to non-compliance with task instructions. Thus, the sample used for analysis included 24 participants (age range 19-32 years, M = 21.50 years, SD = 3.26 years, 14 females). All participants were Italian native speakers and they had normal or corrected-to-normal vision. The experimental protocol is in line with the declaration of Helsinki and with the regulations of the University of Florence which hosted the study.

### Materials

#### Apparatus

Subjects sat in front of a monitor screen, with their heads stabilized with a chin rest. Viewing was binocular. Stimuli were generated with the PsychoPhysics Toolbox routines for MATLAB (MATLAB r2010a, The MathWorks) and presented on a LCD colour monitor (Asus MX239H, 51 x 28 cm placed at 57 cm viewing distance) with a resolution of 1920 x 1080 pixels and a refresh rate of 60 Hz, driven by a Macbook Pro Retina (OS X Yosemite, 10.10.5). All stimuli were shown in white (55 cd/m^2^) against a black background (0.05 cd/m^2^). Two-dimensional eye position and pupil diameter were monitored binocularly with a CRS LiveTrack system (Cambridge Research Systems) at 30 Hz, using an infrared camera mounted below the screen. Pupil diameter measures were transformed from pixels to millimeters after calibrating the tracker with an artificial 4 mm pupil, positioned at the approximate location of the subjects’ left eye. Gaze position data were linearized using a standard 9-point calibration, run prior to each session.

#### Vigilance task

Participants completed a modified version of the computer-based vigilance task [15] already used in previous studies [e.g., 11, 13, 20]. The task consisted of 1020 trials, presented in a fixed order, each lasting 2 s. A white fixation point (0.2 deg diameter) was always shown at screen centre. Each trial presented a pattern of white horizontal (non-target stimuli) or vertical lines (target stimuli) (4.1 x 0.2 deg) randomly distributed across the screen, against a black background. Target stimuli appeared on a total of 30 trials (~3% of all trials) and they were distributed pseudo-randomly, every 26-40 trials; participants were asked to press the space bar whenever a target was detected. Moreover, a white verbal cue (0.88 deg text-height, e.g., “long hair”, “jet lag”) was placed under the fixation spot, in 192 trials (18.8%). The verbal cues were selected from the Italian adaptation of a standardized pool of 800 word-phrases developed by [15] and already used in previous studies (for more details on the adaptation, see 20). Equal numbers of neutral (*n* = 64), positive (*n* = 64), and negative (*n* = 64) cues were presented during the task.

#### Thought questionnaire

After completing the vigilance task, participants were asked to give details of their reported mental contents. For each content, they were asked to indicate: (i) the temporal focus, distinguishing between “past”, “present”, “future”, and “atemporal”, (ii) whether it was general or specific, (iii) the emotional valence of the thought on a 7-point scale (−3 = very unpleasant; 0 = neutral; +3 = very pleasant).

Participants received instructions on how to distinguish the different temporal focus categories. Specifically, they were told that an “atemporal” mental content would refer to any thought with no specific temporal orientation (i.e., “I am very shy”), a “present” mental content would refer to any thought related to something occurring either here and now or in the current period of life, a “past” mental content would refer to any thought related to something occurred prior to begin the task (more or less remote), and a “future” mental content would refer to any thought related to something occurring after the end of the task (more or less distant in the future). Participants were also asked to rate on a 7-point scale their overall level of concentration (1 = not at all concentrated; 7 = fully concentrated) and boredom (1 = not at all; 7 = very bored) experienced during the vigilance task.

### Procedure

Participants were tested individually. When first welcomed into the laboratory, they were briefly introduced to the research project and eye-tracking recording technique, they were informed that they would take part in a study on concentration and its correlates, and they signed a consent form. Then, they received the instructions for the vigilance task: to detect target stimuli (vertical lines) among a large number of non-target stimuli (horizontal lines), by pressing the space-bar after each target stimulus, while keeping their gaze on the fixation point for the whole session. We informed them of the occurrence of verbal cues in some of the trials, which were irrelevant to their task. To justify these, we provided a cover story (which was uncovered after the end of the experiment) – we told participants that the experiment comprised two conditions tested in separate groups of participants, one testing how people focus on the patterns irrespectively of the verbal cues, and the other testing the opposite, focus on the cues irrespectively of the patterns. Finally, participants were told that the task was monotonous and that task-unrelated mental contents (e.g., thoughts, plans, considerations, past events, images, etc.) could pop into their mind spontaneously throughout the task. Any time this happened, they had to press a button on the keyboard (the letter L, marked with a white sticker) to interrupt the presentation and give a short description of the mental content; they would also have to indicate whether it was triggered by internal thoughts, an element in the environment, a cue-word on the screen (if so, they had to specify the word) or no cue at all. These responses were recorded by the experimenter. This initial description should have been sufficient for them to identify the mental content at a later point in time, if necessary. However, if the mental content was private and intimate, participants could label it as “personal” and provide only one relevant word instead of reporting a short description.

After the instructions, participants went through 20 practice trials and finally proceeded to complete two sessions of 510 trials each. At the end of the sessions, subjects completed a questionnaire on their thought experience (see next section) and we asked whether they had speculated about the actual aims of the study (if so, what they had thought) during the task and then they were debriefed and dismissed. The total session lasted approximately 100-120 min.

### Data encoding and analysis

All thoughts recorded by participants were read by two independent judges (first and third authors) and independently coded into three categories: the mental contents could be either task-related interferences (TRIs) or task-unrelated thoughts (TUTs); TUTs were further classified as either external distractions (EDs) or MW episodes. TRIs included all cases where a reference was made to task features or to the participant’s overall performance (i.e., thoughts about the experiment’s duration). EDs included all thoughts focused on task-unrelated sensations, either exteroceptive perceptions (i.e. a noise outside the room) or interoceptive sensations (i.e., bodily sensations). MW episodes included all thoughts that were unrelated to both the task and the sensory environment; however, these thoughts could be triggered by internal or external stimuli (including the verbal cues), which participants identified in their reports. For both categorisations (TRIs *vs*. TUTs, and MW *vs*. EDs), we computed Kappa as inter-rater reliability between the coders and the inter-rater agreement resulted to be very good (*Kappa* = 0.96, *SE* = 0.03 and inter-rater agreement *Kappa* = 0.99, *SE* = 0.01, respectively). Minor disagreements were resolved by discussion.

Next, an off-line analysis examined the eye-tracking output and excluded time-points with unrealistic pupil-size recordings (i.e., values outside the 90^th^ percentile of each 2 sec. long trial) and interpolated the remaining time-points at 20Hz.

To perform the analyses on the pupillometry data, trials were sorted based on their timing relative to a cue later identified as “trigger” or “post-MW report” (i.e., 0, 1, 2 trials after the word, where the trial 0 was the one where the word was shown). We used as “baseline”pupil diameter the average diameter in the reference event (trial “0”). We studied the timecourse of pupil diameter over trials after subtracting this baseline. Since we were interested in the timecourse of MW episodes (0-2, trials after the trigger or 1-3 trials before the self-interruption), we selected only MW events that met three criteria: the latency between the verbal cue and the self-interruption was at least 6 seconds (three trials); in this period, only non-target horizontal lines were shown (to avoid the confounding effects of cue-words and targets on pupil diameter) and reliable pupil recordings were collected.

Statistical analyses relied on a linear-mixed model approach, motivated by the considerable sample size variability across subjects. In this approach, individual trials from all subjects are compared with a model comprising both the effect of experimental variables (“fixed effects”) and the variability across participants (“random effects”). Random effects were coded by allowing subject-by-subject variations of the intercept of the model. In all cases, the dependent variable is “baseline corrected pupil diameter”, which we obtained by averaging pupil diameter in a pre-specified temporal window of each trial and subtracting the average pupil diameter in a reference temporal window. Please refer to the results section for specific definitions of the temporal windows for averaging and baseline-subtractions. We used standard MATLAB functions provided with the Statistics and Machine Learning Toolbox (R2015b, The MathWorks). Specifically, the function “fitlme(data, model)” fits the linear-mixed model to the data, yielding an object “lme” with associated method “ANOVA” that returns *F* statistics with associated degrees of freedom, and *P* values for each of the fixed effect terms and “CoefTest” for post-hoc comparisons.

## Results

### Performance on vigilance task

Performance on the vigilance task was near-perfect for all participants. Out of 30 targets, there were 0.46 (*SD* = 0.66) misses and 0.17 (*SD* = 0.38) false alarms. The mean reaction time associated with correct detections was 767.13 msec. (*SD* = 124.42 msec.). The mean level of concentration experienced during the task was 4.92 (*SD* = 1.06) and the mean level of boredom was 3.04 (*SD* = 1.57).

### Frequency and properties of MW reports

Participants reported a total of 400 mental contents (*M* = 16.67, *SD* = 16.83, range 1-63). Out of the all mental contents, 27 reports (6.75%) were classed as TRI reports (*M* = 1.13, *SD* = 1.54, range 0-6), and 373 reports (93.25%) were classed as TUT reports (*M* = 15.54, *SD* = 16.59, range 1-61). Out of all the TUTs, 40 reports (10.72%) were classed as external distractions (*M* = 1.67, *SD* = 1.93, range 0-6) and 333 reports (89.28%) were classed as MW (*M* = 13.88, *SD* = 15.48, range 0-58). One out of 333 MW reports was excluded from the analysis of pupil diameter because of inaccurate recording of time of MW interruption and report.

Out of 333 MW episodes, 67.57% were reported as triggered by a verbal cue previously shown on-screen, 3.90%, by internal thoughts, 4.50% by environmental triggers and 18.32% by no trigger. The remaining 19 MW episodes were indicated by participants as elicited by multiple cue-words (i.e., more than one specific cue-word; n =7), by unknown cue-word(s) (i.e., participants reported that they did not remember which cue-word triggered the MW episode; n = 8), or by multiple triggers (e.g., participants reported that their thoughts were elicited by both a cue-word presented onscreen and a sound occurring outside the room; n = 4).

Finally, out of all the MW episodes triggered by the verbal cues, 26.67% were triggered by neutral cue-words, 35.11% by positive cue-words and 38.22% by negative cue-words. Thus, the vast majority (73.33%) of these reports were elicited by verbal cues with emotional valence.

At the end of the vigilance task, participants coded each of their reported thoughts as past-oriented, present-oriented, future-oriented or atemporal thoughts. Out of the all MW episodes, 133 episodes (39.94%) were classed as past-oriented thoughts, 21 episodes (6.31%) were classed as present-oriented thoughts, 53 episodes (15.91%) were classed as future-oriented thoughts, and 126 episodes (37.84%) were classed as atemporal thoughts. Moreover, out of all the MW episodes, 167 episodes (50.15%) were classed as specific, whereas 166 episodes (49.85%) were classed as general. Finally, 96 episodes (28.83%) were rated as neutral, 118 episodes (35.43%) were rated as negative, and 119 (35.74%) were rated as positive.

### Latency data

For each MW episode indicated by participants as triggered by a cue-word, we computed the time-interval between a verbal cue and the task-interruption where a MW episode was reported as triggered by said verbal cue (see also studies on involuntary memories for a similar procedure to obtain retrieval times; e.g., 15, 20)

Out of the 225 MW episodes triggered by a verbal cue, 5 were identified as outliers (more than 2.5 standard deviations away from the mean latency). The mean latency of the remaining MW episodes was 7.291s (*SD* = 5.807s), which can be interpreted as the average duration of the MW episode (from its onset, marked by the trigger presentation, to its end, marked by the self-interruption of the task). The distribution of latency data is reported in Fig 1.

**Fig 1.**
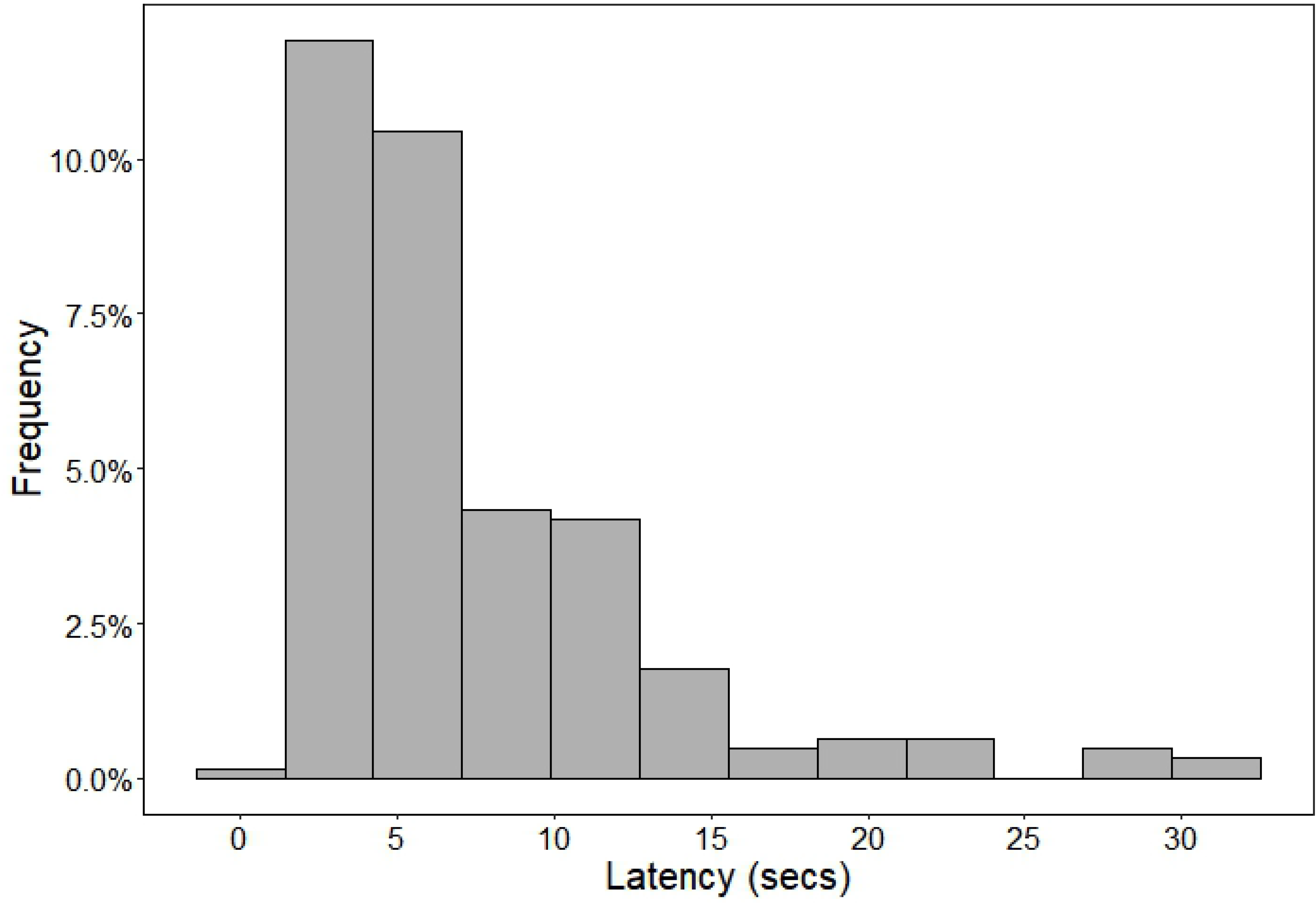
Distribution of latency data. Distribution of the time-intervals between a verbal cue and a MW report.

We analyzed whether and how the phenomenological characteristics of MW episodes (temporal focus, specificity and emotional valence) affected their durations by means of a multilevel (or hierarchical) analysis. We specified random-intercept multilevel models to test for associations of the factors: temporal focus (past, present, future, atemporal), specificity (general, specific), valence (negative, neutral, positive). With regard to the valence of thoughts, participants rated each episode on a 7-point Likert scale (−3 = very negative; 0 = neutral; +3= very positive) and we used these ratings to classify the episode as negative (scored between −3 and −1), neutral (scored 0) or positive (scored between +1 and +3).

Given that the MW episode durations were substantially skewed and kurtotic, we conducted the analysis after log transformation of the data.

We found no significant effect of temporal focus, *F*(3, 201.24) =0.33, *p* = 0.806 (past Estimated Marginal Mean = 3.83, 95% Confidence Interval [CI]: 3.67-3.98; present Estimated Marginal Mean = 3.77, 95% CI: 3.57-3.96; future Estimated Marginal Mean = 3.82, 95% CI: 3.65-3.99; atemporal Estimated Marginal Mean = 3.80, 95% CI: 3.64-3.95). We also found no significant duration difference between episodes associated with different emotional valences, *F*(2, 202.64) = 0.41, *p* = 0.665 (neutral Estimated Marginal Mean = 3.79, 95% Confidence Interval [CI]: 3.63-3.95; negative Estimated Marginal Mean = 3.82, 95% CI: 3.67-3.97; positive Estimated Marginal Mean = 3.82, 95% CI: 3.67-3.98), and no significant difference between specific and general episodes, *F*(1, 205.13) = 3.47, *p* = 0.064 (specific Estimated Marginal Mean = 3.84, 95% Confidence Interval [CI]: 3.69-3.98; general Estimated Marginal Mean = 3.78, 95% CI: 3.63-3.93).

### Pupillometric correlates of mind wandering

In the first analysis of pupillometry data, we analysed the timecourse of pupil diameter over two trials after a trigger (a verbal cue presented before a MW event and identified by the participant as the trigger of said event, n=107) or a non-trigger (a verbal cue presented after a MW report, hence unlikely to be trigger of further distractions, reported or non-reported, n=226).

Fig 2 shows the traces, aligned to the average pupil diameter during the verbal cue presentation (baseline window defined as the first half of the verbal-cue trial). Verbal cues (white text on black background) evoked strong pupillary constriction (the pupillary light response), which recovers over several seconds producing a progressive pupil dilation. This pupil dilation is more pronounced for trigger cues than non-trigger cues.

**Fig 2.**
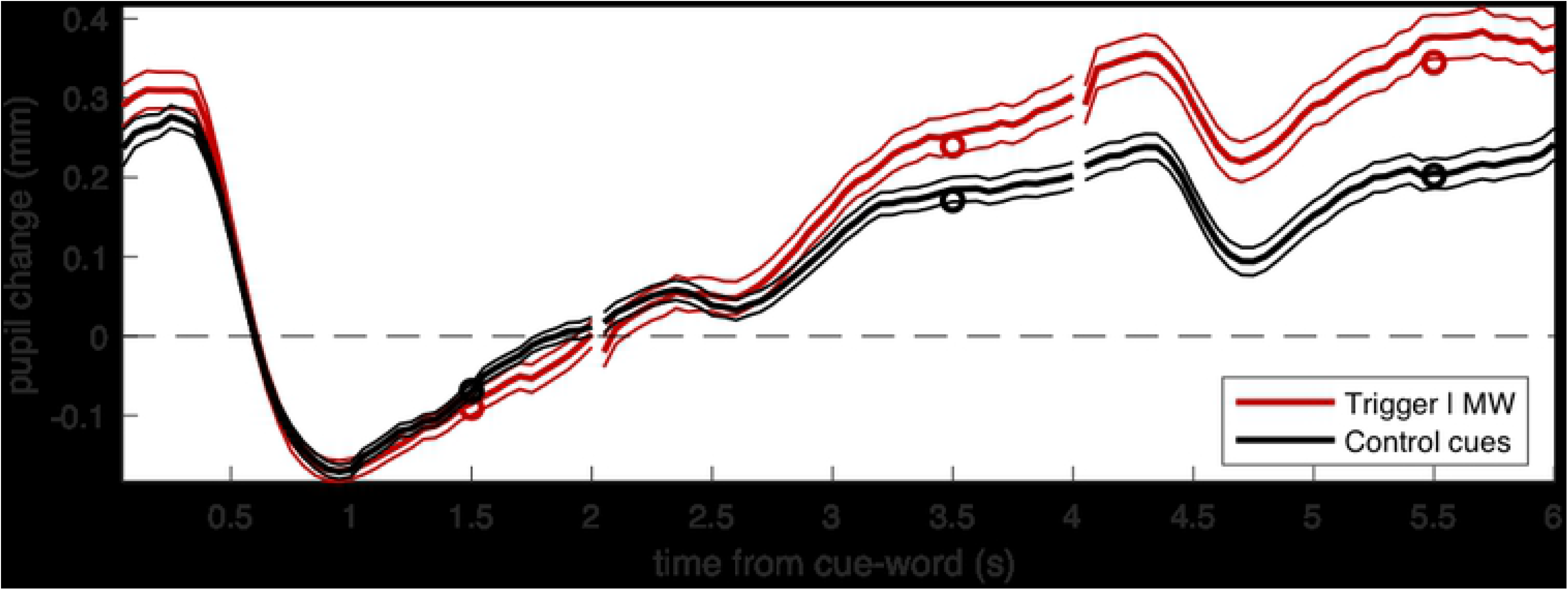
Pupil traces aligned to the average pupil diameter during the presentation of the verbal cue. Thick lines give the average across all trials, thin lines show the s.e. and circles show the average values entered in the LMM analysis (average over the second half of each trial). The two traces give the average pupil diameter following a verbal cue that was later reported as triggering a MW episode, or a verbal cue presented during a supposedly on-task period, i.e. the period immediately following the resuming of the task after a self-caught interruption and MW report.

In order to statistically assess this effect, we summarised pupil traces by taking the average pupil diameter in the last second of each trial (the farthest from the baseline window). These values were entered in a Linear-Mixed Model analysis, with two fixed-factors: type of cue-word (trigger and non-trigger) and time from the cue (coded as number of trials: 1^st^ trial or 2^nd^ trial after the cue-word), plus the random effect of subjects modelled as a variable intercept of the model. This revealed a significant interaction between the two fixed factors (F(1,995) = 23.48584, p < 0.00001), which we further analysed with a series of post-hoc tests. These showed that cue-type has a significant effect on both trials following the cue, the first (F(1,331) = 9.15235, p = 0.00268) and the second (F(1,331) = 22.41800, p < 0.00001). These findings were confirmed when we used as control-cues a subset of word-cues with emotional content (N=156) – compared against the same N=107 trigger-cues which, for the most part, had emotional content. The Linear-Mixed model analysis revealed a significant interaction between type of verbal cue (trigger, and emotional non-trigger) and time from the cue (F(1,785) = 19.01058, p = 0.00001), and again indicated a large effect of cue-type on both trials following the cue, the first (F(1,261) = 10.38556, p = 0.00143) and the second (F(1,261) = 19.99679, p = 0.00001).

Following [21] we complemented this with a second analysis, aligning traces to the MW reports (Fig 3) or to “control trials” which we artificially set at 5 trials following every selfinterruption – so close to an interruption that the subjects were very unlikely to have leaped back into a mind wandering state.

**Fig 3.**
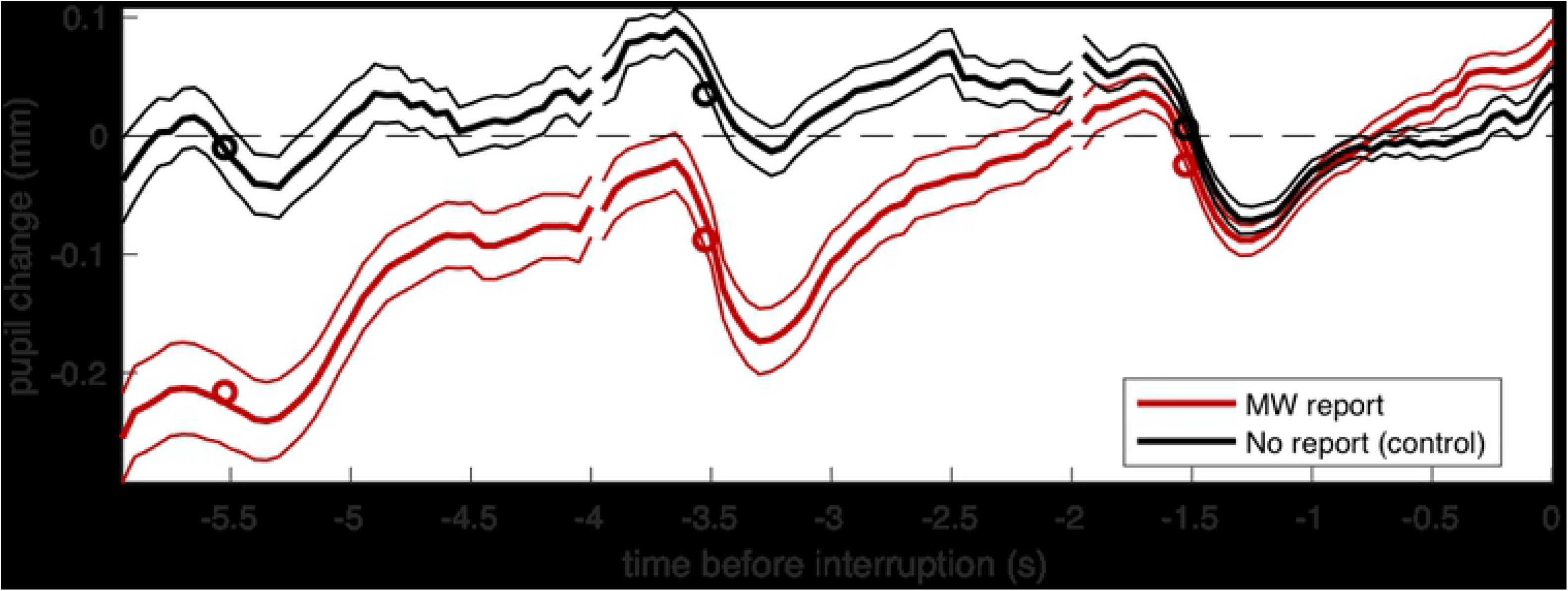
Pupil traces aligned to the average pupil diameter on the last trial before the MW report or the control trial. Thick lines give the average across all trials, thin lines show the s.e. and circles show the average values entered the LMM analysis (average over the second half of each trial).

Pupil diameter preceding these control trials can also be seen as the timecourse that pupil diameter should regularly show in this experiment, without the intrusion of a mind wandering episode. Also, in this case, there was a relative pupil dilation preceding the MW report. To show this statistically, we aligned traces to the average pupil diameter on the last trial before the MW report or the control trial, then assessed pupil diameter on each trial as the mean pupil diameter in the first second of the trial (the farthest from the reference). This left 89 MW reports and 102 control trials. The Linear-Mixed model analysis revealed a significant interaction between the fixed-factors type of trial (MW/control) and time (F(1,569) = 18.22662, p = 0.00002). At three trials preceding the reference trial, pupil diameter leading to a MW report could be clearly differentiated from the usual pupil diameter during the experiment (F(1,189) = 25.47720, p < 0.00001). This confirms that there is pupil dilation leading up to a MW report.

## Discussion

Several recent studies have shown that MW is a cue-dependent phenomenon, triggered by both internal and external events. Specifically, there is evidence that both task-irrelevant and task-relevant external stimuli [7–11, 13] might act as triggers for MW episodes. In a recent study, Pelagatti et al. [21] found a significantly larger pupil dilation following cue-words reported by participants as the trigger of MW compared to non-trigger words (with similar emotional content), and the pupil dilation appeared to increase over time. In the study by Pelagatti et al. [21] MW was assessed by using the probe-catching method. In the present study, we globally replicated these findings using the self-catching method, which *“provides a straightforward assessment of the number of mind wandering episodes that reached meta-awareness”* [24, p.322], thereby allowing investigation of the pupil correlates of MW that an individual becomes aware of (aware MW).

Our results on the increased pupil diameter during MW are in line with some previous findings [e.g. 21, 29; see also 30, for larger pupil diameters in participants who retrospectively reported more MW compared to those who reported less MW). However, other studies have reported the opposite finding: reduced pupil dilation during MW [e.g. 27, 31–33]. The control of pupil diameter is affected by a variety of factors [e.g. 34–43], and ultimately reflects the balance between the parasympathetic and sympathetic systems (reflecting the level of arousal, e.g. as linked with cognitive and emotional load, see 23, 41, 44). When light level is constant, we can expect pupil dilation every time the cognitive or emotional load is increased [e.g., 23]. However, given any two tasks, it is not always obvious which of the two is associated with higher load. In the MW literature, pupil diameter is compared between two conditions: epochs when the participant is focused on the task, and seldom interspersed with mind-wandering events. Are the latter events associated with more or less load than on-task epochs? And, consequently, should we expect pupil dilation or constriction during a MW episode? We believe that the answers to these questions will always depend on the exact characteristics of the main task and of the MW episodes. In this sense, we agree with Konishi et al. (32; see also 31, 45), suggesting that the content of MW and the context in which it emerges are determinant for its relation to other neurocognitive variables (including pupil size). Therefore, it is impossible, a priori, to predict the direction of pupil modulations for a new experimental setting. This does not reduce the importance of pupillometry, which emerges as a reliable marker of MW within each experimental context. In our case, the presence of pupil dilation during MW episodes may be interpreted considering two aspects: the monotonous and undemanding main task (more repetitive and less challenging than other tasks used in previous research on pupillometry and mw) and the presence of cue-words, most often held responsible for the occurrence of the MW episodes, which might have charged the latter with particularly high emotional/cognitive load. Moreover, the presence of cue-words also gave us the opportunity to estimate the latency of MW episodes that these had triggered, by computing the time-interval between the presentation of triggering cue-words and the self-reports of MW. The latency of a MW episode could be considered as the time needed for the arising of thought and for becoming aware of it in order to report it. We found high variability in the duration of MW, with a median latency of approximately 6 sec and a percentage of 41% MW episodes lasting 5 sec or less. These findings are in line with the ones reported by Klinger [4], who trained participants to estimate the duration of their thoughts and found a median estimate of thought segment duration of 5 sec.

Our results, combining behavioral and pupillometric measures, have implications at both theoretical and methodological level. So far, most research on MW has adopted a content-based approach to MW, focusing on the property of task-relatedness to distinguish MW from other attentional states; however, the dynamic aspects of MW, which also includes the temporal duration, have not been thoroughly considered. Most MW studies have used fixed timewindows to compare on-task and MW episodes; however, the duration of such time-windows is highly variable across studies [e.g., 10 sec. in 26, 46; 6.5 sec. in 47; 3 to 8 sec. in 48; 5 sec. in 49, 50; 4.8 sec. in 51] thereby testifying the scarcity of evidence on the temporal unfolding of MW. Moreover, by using a fixed time-window, identical for all episodes and participants, these studies have implicitly assumed that the duration of MW episodes is relatively constant.

However, both our results and the ones reported by Klinger [4] clearly show that the duration of MW is highly variable, both within subjects (across different MW episodes) and between subjects. These findings should stimulate research into the factors that might explain and predict this variability and help our understanding of the processes that lead to the variable timing of MW.

In conclusion, classic experiments on MW have left many of its key properties, and particularly its temporal properties, unexplored. Future research will need new objective tools to clarify these aspects; we submit that our pupillometry paradigm can contribute to this quest, providing an objective and reliable marker of the onset and temporal unfolding of MW episodes.

## References

1. Smallwood, J O’Connor RC, Sudbery MV, Obonsawin M. Mind-wandering and dysphoria. Cogn Emot. 2007 21(4): 816–842. doi: 10.1080/02699930600911531

2. Smallwood J, Schooler JW. The science of mind wandering: Empirically navigating the stream of consciousness. Annu Rev Psychol. 2015; 66: 487–518. doi: 10.1146/annurev-psych-010814-015331

3. Smallwood J. Distinguishing how from why the mind wanders: A process-occurrence framework for self-generated mental activity. Psychol Bull. 2013; 139 (5): 519–535. doi: 10.1037/a0030010

4. Klinger E. Modes of normal conscious flow. In: Pope KS, Singer JL, editors, The stream of consciousness. Emotions, personality, and psychotherapy. Boston, MA: Springer; 1978. pp. 225–258. doi:10.1007/978-1-4684-2466-9_9

5. Klinger E. Goal commitments and the content of thoughts and dreams: Basic principles. Front Psychol. 2013; 4: 415. doi: 10.3389/fpsyg.2013.00415

6. Klinger E, Marchetti I, Koster E. Spontaneous thought and goal pursuit: From functions such as planning to dysfunctions such as rumination. In Fox KCR, Christoff K, editors. The Oxford handbook of spontaneous thought: Mind wandering, creativity, dreaming, and clinical disorders. Oxford, UK: Oxford University Press. 2018. pp. 215–232. doi:10.1093/oxfordhb/9780190464745.013.24

7. Jordão M, Pinho MS, St. Jacques PL. Inducing spontaneous future thoughts in young and older adults by priming future-oriented personal goals. Psychol Res. 2019; 83 (4): 710–726. doi: 10.1007/s00426-019-01146-w

8. Maillet D, Schacter DL. When the mind wanders: Distinguishing stimulus-dependent from stimulus-independent thoughts during incidental encoding in young and older adults. Psychol Aging. 2016; 31 (4): 370–379. doi: 10.1037/pag0000099

9. Maillet D, Seli P, Schacter D. Mind-wandering and task stimuli: Stimulus-dependent thoughts influence performance on memory tasks and are more often past versus future-oriented. Conscious Cogn 2017; 52: 55–67. doi: 10.1016/j.concog.2017.04.014

10. McVay JC, Kane MJ. Dispatching the wandering mind? Toward a laboratory method for cuing “spontaneous” off-task thought. Front Psychol. 2013; 4: 570. doi: 10.3389/fpsyg.2013.00570

11. Plimpton B, Patel P, Kvavilashvili L. Role of triggers and dysphoria in mind-wandering about past, present and future: A laboratory study. Conscious Cogn, 2015; 33: 261–276. doi: 10.1016/j.concog.2015.01.014

12. Song X, Wang X. Mind wandering in Chinese daily lives – An experience sampling study. PLoS ONE, 2012; 7(9), e44423. doi: 10.1371/journal.pone.0044423

13. Vannucci M, Pelagatti C, Marchetti I. Manipulating cues in mind wandering: Verbal cues affect the frequency and the temporal focus of mind wandering. Conscious Cogn. 2017; 53: 61–69. doi: 10.1016/j.concog.2017.06.004

14. Antrobus JS. Information theory and stimulus-independent thought. Br. J. Psychol. 1968; 59: 423–430. doi: 10.1111/j.2044-8295.1968.tb01157.x

15. Schlagman S, Kvavilashvili L. Involuntary autobiographical memories in and outside the laboratory: How different are they from voluntary autobiographical memories? Mem Cognit. 2008; 36 (5): 920–932. doi: 10.3758/MC.36.5.920

16. Barzykowski K, Niedźwieńska A. The effects of instruction on the frequency and characteristics of involuntary autobiographical memories. PLoS One. 2016; 11 (6): e0157121. doi: 10.1371/journal.pone.0157121

17. Barzykowski K, Staugaard SR. Does retrieval intentionality really matter? Similarities and differences between involuntary memories and directly and generatively retrieved voluntary memories. Br. J. Psychol. 2015; 107(3): 519–536. doi: 10.1111/bjop.12160

18. Vannucci M, Batool I, Pelagatti C, Mazzoni G. Modifying the frequency and characteristics of involuntary autobiographical memories. PLoS One. 2014; 9(4), e89582. doi: 10.1371/journal.pone.0089582

19. Vannucci M, Pelagatti C, Chiorri C, Mazzoni G. Visual object imagery and autobiographical memory: object imagers are better art remembering their personal past. Memory. 2016; 24 (4): 455–470. doi: 10.1080/09658211.2015.1018277

20. Vannucci M, Pelagatti C, Hanczakowski M, Mazzoni G, Rossi Paccani C. Why are we not flooded by involuntary autobiographical memories? Few cues are more effective than many. Psychol Res. 2015; 79(6): 1077–1085. doi: 10.1007/s00426-014-0632-y

21. Pelagatti C, Binda P, Vannucci M. Tracking the dynamics of mind wandering: Insight from pupillometry. J Cogn. 2018; 1(1): 1–12. doi: 10.5334/joc.41

22. Hess EH, Polt JM. Pupil size as related to interest value of visual stimuli. Science. 1960; 132 (3423): 349–350. doi: 10.1126/science.132.3423.349

23. Kahneman D, Beatty J. Pupil diameter and load on memory. Science. 1966; 154(3756), 1583–1585. doi: 10.1126/science.154.3756.1583

24. Schooler JW, Smallwood J, Christoff K, Handy TC, Reichle ED, Sayette MA. Meta-awareness, perceptual decoupling and the wandering mind. Trends Cogn Sci. 2011; 15 (7): 319–326. doi: 10.1016/j.tics.2011.05.006

25. Smallwood J, McSpadden M, Schooler JW. When attention matters: The curious incident of the wandering mind. Mem Cognit. 2008; 36 (6): 1144–1150. doi: 10.3758/MC.36.6.1144

26. Christoff K, Gordon AM, Smallwood J, Smith R, Schooler JW. Experience sampling during fMRI reveals default network and executive system contributions to mind wandering. Proc Natl Acad Sci USA. 2009; 106 (21): 8719–8724. doi: 10.1073/pnas.0900234106

27. Grandchamp R, Braboszcz C, Delorme A. Oculometric variations during mind wandering. Front Psychol. 2014; 5: 31. doi: 10.3389/fpsyg.2014.00031

28. Pope KS. The flow of consciousness. Ph.D. dissertation, Yale University. 1977.

29. Franklin MS, Broadway JM, Mrazek MD, Smallwood J, Schooler JW. Window to the wandering mind: Pupillometry of spontaneous thought while reading. Q J Exp Psychol (Hove) 2013; 66 (12): 2289–2294. doi: 10.1080/17470218.2013.858170

30. Smallwood J, Brown KS, Tipper C, Giesbrecht B, Franklin MS, Mrazek MD, Carlson JM, Schooler JW. Pupillometric evidence for the decoupling of attention from perceptual input during offline thought. PLoS One. 2011; 6(3): e18298. doi: 10.1371/journal.pone.0018298

31. Gouraud J, Delorme A, Berberian B. Influence of automation on mind wandering frequency in sustained attention. Conscious Cognit. 2018; 66: 54–64. doi: 10.1016/j.concog.2018.09.012.

32. Konishi M, Brown K, Battaglini L, Smallwood J. When attention wanders: Pupillometric signatures of fluctuations in external attention. Cognition. 2017; 168: 16–26. doi:10.1016/j.cognition.2017.06.006

33. Unsworth N, Robison MK. Pupillary correlates of lapses of sustained attention. Cogn Affect Behav Neurosci. 2016; 16(4): 601–615. doi: 10.3758/s13415-016-0417-4

34. Binda P, Gamlin PD. Renewed Attention on the Pupil Light Reflex. Trends Neurosci. 2017; 40: 455–457. doi: 10.1016/j.tins.2017.06.007.

35. Binda P, Murray SO. Spatial attention increases the pupillary response to light changes. J Vis. 2015; 15 (2): 1. doi: 10.1167/15.2.1.

36. Binda P, Murray SO. (2015). Keeping a large-pupilled eye on high-level visual processing. Trends Cogn Sci. 2015; 19: 1–3. doi: 10.1016/j.tics.2014.11.002.

37. Binda P, Pereverzeva M, Murray SO. Attention to bright surfaces enhances the pupillary light reflex. J Neurosci. 2013a; 33: 2199–2204. doi: 10.1523/JNEUROSCI.3440-12.2013.

38. Binda P, Pereverzeva M, Murray SO. Pupil constrictions to photographs of the sun. J Vis. 2013b; 13: 8. doi: 10.1167/13.6.8.

39. Binda P, Pereverzeva M, Murray SO. Pupil size reflects the focus of feature-based attention. J Neurophysiol. 112(12): 3046–3052. doi: 10.1152/jn.00502.2014.

40. Laeng B, Endestad T. Bright illusions reduce the eye’s pupil. Proc Natl Acad Sci U S A. 2012; 109: 2162–2167. doi: 10.1073/pnas.1118298109

41. Laeng B, Sirois S, Gredebäck G. Pupillometry: A Window to the Preconscious? Perspect Psychol Sci. 2012; 7 (1): 18–27. doi: 10.1177/1745691611427305

42. Mathôt S, Van der Linden L, Grainger J, Vitu F. The pupillary light response reveals the focus of covert visual attention. PLoS ONE. 2013; 8(10), e78168. doi: 10.1371/journal.pone.0078168

43. Turi M, Burr DC, Binda P. Pupillometry reveals perceptual differences that are tightly linked to autistic traits in typical adults. Elife 2018; 7. doi: 10.7554/eLife.32399.

44. Einhauser W, Stout J, Koch C, Carter O. Pupil dilation reflects perceptual selection and predicts subsequent stability in perceptual rivalry. Proc Natl Acad Sci U S A. 2009; 105(5): 1704–9. doi: 10.1073/pnas.0707727105

45. Jubera-Garcia E, Gevers W, Van Opstal F. Influence of content and intensity of thought on behavioral and pupil changes during active mind-wandering, off-focus, and on-task states. Attention Perception and Psychophysics. 2019; Epub ahead of print. doi: 10.3758/s13414-019-01865-7

46. Franklin MS, Smallwood J, Schooler J. Catching the mind in flight:_Using behavioral indices to detect mindless reading in real time. Psychon Bull Rev 2011; 18: 992–997. doi:10.3758/s13423-011-0109-6

47. Seli P, Cheyne JA, Smilek, D. Wandering minds and wavering rhythms:linking mind wandering and behavioral variability. J Exp Psychol Hum Percept Perform 2013; 39(1):1–5. doi: 10.1037/a0030954

48. Frank DJ, Nara B, Zavagnin M, Touron DR, Kane MJ. Validating older adults’ reports of less mind-wandering: An examination of eye-movements and dispositional influences. Psychol Aging. 2015; 30: 266–278. doi:10.1037/pag0000031

49. Uzzaman S, Joordens S. The eyes know what you are thinking: eye movements as an objective measure of mind wandering. Conscious Cogn 2011; 20(4): 1882–6. doi: 10.1016/j.concog.2011.09.010

50. Smilek D, Carriere JSA, Cheyne A. Out of mind, out of sight: Eye_blinking as indicator and embodiment of mind wandering. Psychol Science; 2010; 21(6): 786–789. doi:10.1177/0956797610368063

51. McVay JC, Kane MJ. Conducting the train of thought: Working memory capacity, goal neglect, and mind wandering in an executive-control task. J Exp Psychol: Lear, Mem Cogn, 2009; 35: 196–204. doi:10.1037/a0014104

